# Rac1 inhibition prevents axonal cytoskeleton dysfunction in Transthyretin Amyloid Polyneuropathy

**DOI:** 10.1101/2024.11.27.625664

**Authors:** Joana Magalhães, Vítor Dias, Jessica Eira, Guilherme Nóvoa, Marina I. Oliveira da Silva, Yu Yu Kan, Andreia Dias, Estefânia Carvalho, Mariana Santos, Ana I Seixas, Teresa Coelho, Ricardo Taipa, Joana Paes de Faria, Carolina Lemos, Sung-Tsang Hsieh, Márcia A. Liz

## Abstract

Hereditary transthyretin amyloidosis with polyneuropathy (ATTRv-PN) is characterized by the deposition of amyloidogenic TTR, particularly in dorsal root ganglia (DRG) and peripheral nerve axons, resulting in sensorimotor axonopathy with autonomic dysfunction. Here, we investigated cytoskeleton alterations in peripheral axons from an ATTRv-PN mouse model, the hTTRA97S knock-in mice. Proteomics of hTTRA97S sural nerves revealed dysregulation of actin-related proteins. hTTRA97S DRG neurons presented a defective actin distribution in growth cones along with a reduction in axonal actin trails, and an associated impairment in the pool of pre-synaptic vesicles. Microtubule dynamics and axonal transport abnormalities were also observed in mutant axons. Importantly, cytoskeletal defects in hTTRA97S neurons preceded axonal degeneration and were mediated by Rac1 hyperactivation in DRG neurites and sciatic nerves of pre-symptomatic mice. Rac1 inhibition rescued cytoskeleton alterations, preventing degeneration. Remarkably, in ATTRv-PN patients with late-onset disease, a variant in RACGAP1, encoding a Rac1 inactivator, supported the neuroprotective role of Rac1 inhibition. Our findings demonstrate that cytoskeletal defects precede axonal degeneration in ATTRv-PN and highlight Rac1 as a promising therapeutic target.

## Introduction

Hereditary transthyretin amyloidosis with polyneuropathy (ATTRv-PN) is a fatal autosomal dominant neurodegenerative disease, characterized by the deposition of oligomers, aggregates, and amyloid fibrils of mutated transthyretin (TTR), particularly in the peripheral nervous system (PNS), leading to a distal axonopathy [1]. Age of disease-onset in ATTRv-PN presents great variabilities. Early-onset ATTRv cases (less than 50 years of age) are characterized by small-fiber-predominant axonal degeneration, leading to sensory dissociation, namely abnormal sensation symptoms, primarily due to amyloid fibril deposition around nerve fibers. In contrast, late-onset cases (more than 50 years of age) involve both small and large fiber damage with loss of all sensory modalities, despite less amyloid deposition relative to the severity of nerve fiber loss [2–4]. These observations suggest the existence of mechanisms other than direct damage caused by amyloid fibers.

The available therapeutic options for ATTRv-PN comprise liver transplantation (the major organ of TTR synthesis), which slows disease progression in young-onset patients [5, 6] although it is highly invasive. Other therapies available include the use of TTR stabilizers and silencing *TTR* gene expression [7–10]. Besides the fact that the above therapies can only be applied after disease onset, none of the available strategies targets the ongoing axonal degeneration. As such, the need to further characterize cellular and molecular pathways involved in ATTRv-PN-related axonopathy persists.

Abnormalities in cytoskeletal organization have been reported in several neurodegenerative disorders [11]. In the case of ATTRv-PN, the dying-back pattern of axonal degeneration suggests an initial disturbance of the distal cytoskeleton as a consequence of TTR deposition. Using an ATTRv-PN *Drosophila* model in which the amyloidogenic mutant ATTRV30M was expressed in photoreceptor cells resulting in roughening of the eye we reported that actin cytoskeleton defects are involved in ATTRv-PN pathogenesis [12]. Specifically, ATTRV30M-expressing flies exhibited defective axonal projections of photoreceptor neurons, characterized by more compact growth cones lacking the spread distribution of filopodia and lamellipodia actin structures [12]. A genetic screen further revealed that reducing levels of Rac1 and Cdc42, members of the Rho GTPase family, ameliorated the TTR-induced rough eye phenotype while decreasing RhoA levels worsened it. Rac1 silencing also rescued the axonal and growth cone defects in ATTRV30M photoreceptors [12].

Here we explored the role of cytoskeletal dysregulation, and Rac1 involvement in ATTRv-PN using a mouse model of the disease, the hTTRA97S mice [13]. These mice replicate the early degeneration of sensory nerves observed in ATTRv-PN [13]. The hTTRA97S mouse is a knock-in model generated by replacing one allele of the mouse *Ttr* locus with human *TTR* bearing the A97S mutation (hTTRA97S) or, in the case of control mice, with human WT *TTR* (hTTRWT) [13]. By approximately 24 months of age, the hTTRA97S mice present TTR deposition in peripheral nerves, decreased density of myelinated axons in the sural nerve (a purely sensory nerve), loss of intraepidermal nerve fibers in the skin, and mechanical allodynia [13]. Besides being a powerful model to test new therapeutic drugs, hTTRA97S mice are a valuable tool for dissecting the intracellular alterations occurring as a consequence of TTR deposition.

Our findings reveal that Rac1-mediated cytoskeleton dysfunction, including disruption of the actin-vesicle network, and defective microtubule dynamics and axonal transport, precedes axonal degeneration in ATTRv- PN. Notably, we demonstrate that Rac1 inhibition prevents neurodegeneration of hTTRA97S peripheral neurons. Importantly, we detected a genetic variant leading to the upregulated expression of the *RACGAP1* gene, which encodes for the specific Rac1 inactivator Rac GTPase-activating protein 1 (RacGAP1) [14], in late-onset patients, strongly suggesting that a decrease in Rac1 activity is neuroprotective. Collectively, we suggest Rac1 as a novel therapeutic target for ATTRv-PN.

## Results

### Amyloidogenic TTR upregulates actin cytoskeleton molecular pathways in sensory axons

Characterizing the proteome profile of tissues affected by amyloidogenic TTR might elucidate the molecular mechanisms underlying ATTRv-PN pathogenesis. While previous studies have focused on tissues such as the liver, adipose tissue, and salivary glands from ATTR patients [26, 27], there is no available data on gene expression or proteomics profile of disease in sensory neurons. To advance our understanding of axonal dysfunction in ATTRv-PN, we performed proteomic analysis of the sural nerve in 9-month-old hTTR97S mice, a time point in which no axonal degeneration was observed, as evidenced by the unchanged density of myelinated axons in the sural nerve (Fig. 1a,b) and the constant number of intraepidermal nerve fibers in the hind paw pad skin (Fig. 1c,d). Moreover, we found that 9-month-old hTTRA97S mice displayed no defects in sensory performance as determined by the von Frey test (Fig. 1e), previously shown to be impaired in 24- month-old animals [13]. These results confirm that 9 months is a pre-symptomatic stage of the disease, allowing us to characterize early proteomic alterations triggered by hTTRA97S. Additionally, both human and mouse TTR levels were comparable in hTTRA97S and hTTRWT sural nerves (Supplementary Fig. 1a-c).

**Fig. 1.**
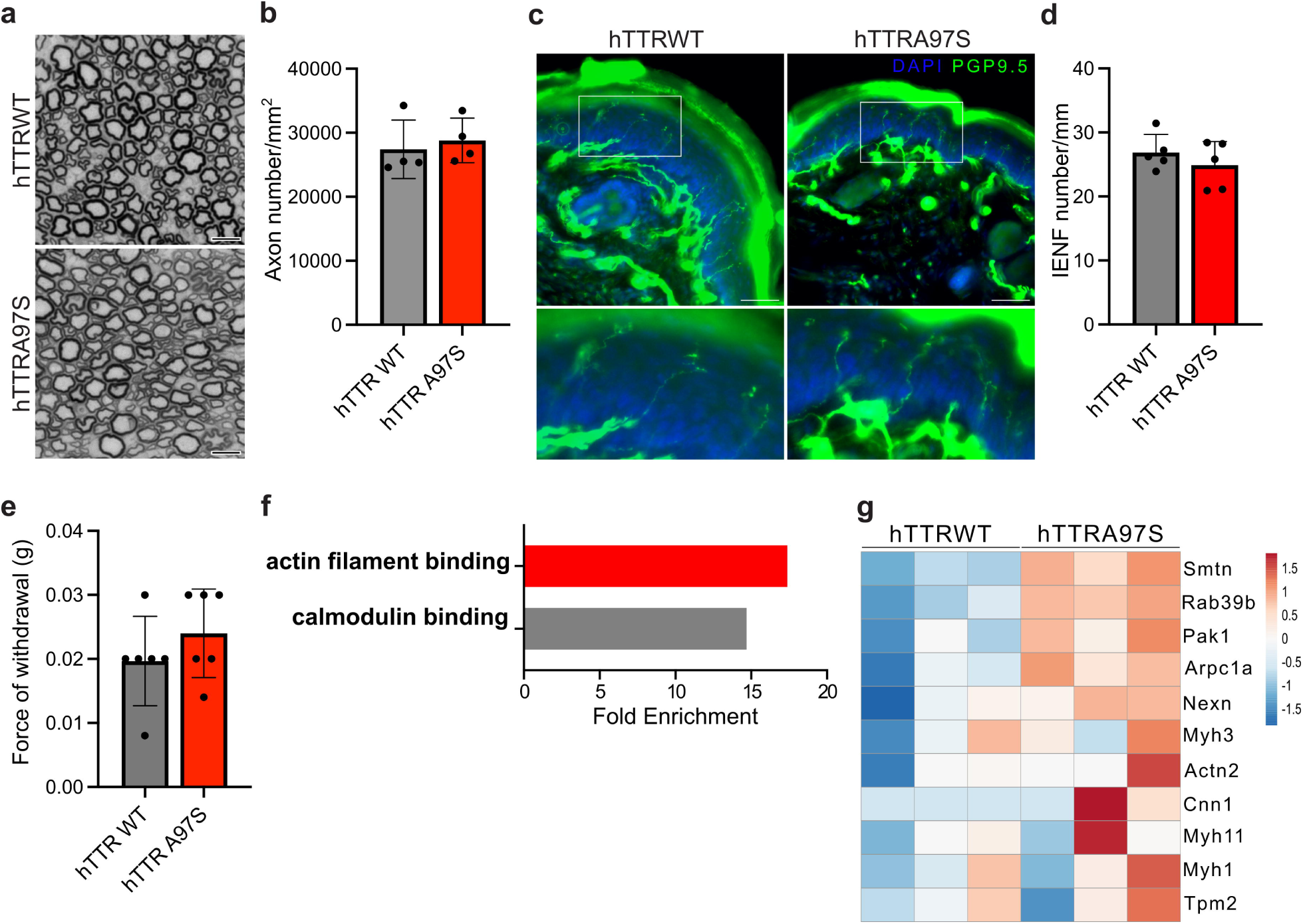
Amyloidogenic TTR upregulates actin cytoskeleton molecular pathways in sensory axons before disease onset. **a,b** Representative images of toluidine blue stained semi-thin sections of hTTRWT and hTTRA97S sural nerves at 9 months of age **(a)** and corresponding density of myelinated axons **(b)**. Scale bar = 5μm. Data represent mean±SD (n=5-6 animals/genotype). **c** Representative images of footpad sections from hTTRWT and hTTRA97S mice immunostained for protein gene product 9.5 (green); DAPI (blue). Scale bar = 50 μm. **d** Quantification of intraepidermal nerve fiber (IENF) density relative to c. Results presented as mean ± SD (n= 5 animals/genotype). **e** von Frey test in 9-month-old hTTRWT and hTTRA97S. Data represent mean±SD (n=6 animals/genotype). **f** Fold enrichment of gene ontology categories for molecular function of upregulated proteins in hTTRA97S sural nerves. Gene ontology analyses were performed using PANTHER 19.0. **g** Heat map of actin-related proteins differentially expressed between 9-month-old hTTRWT and hTTRA97S mice.

Our proteomic analysis revealed 245 proteins that were differentially regulated in hTTRA97S nerves compared to hTTRWT controls (Supplementary Table 2). Gene ontology (GO) analysis of the 46 proteins with increased levels in mutant nerves revealed alterations in molecular pathways primarily related to actin filament binding and calmodulin binding, with 11 and 6 dysregulated proteins, respectively (Fig. 1f). The enrichment analysis of downregulated proteins exposed a scattered scenario, with 199 proteins differentially expressed revealing alterations in a wide range of molecular pathways. The significant enrichment of actin-binding proteins (Fig. 1g) among the differentially regulated proteins in hTTRA97S sural nerves supports our previous findings linking axonal cytoskeleton and ATTRv-PN neurodegeneration in a *Drosophila* model [12]. Building on the integration of these proteomic data with our published work, we proceeded to analyse actin organization in hTTRA97S neurons.

### hTTRA97S sensory neurites have a defective F-actin network preceding axonal degeneration

To uncover actin cytoskeleton defects in the ATTRv-PN mouse model, we resorted to DRG neuron cultures to analyse sensory neurites in the technical impossibility of investigating actin dynamics in sural nerve axons. We initially assessed whether hTTRA97S DRG neurites presented actin cytoskeleton alterations at the growth cone, similarly to what we have observed in photoreceptor axons from ATTRv-PN flies [12]. Although only present in growing neurons, the growth cone is a suitable cellular structure to study neuronal cytoskeleton rearrangements, as both microtubule and actin are stereotypically organized within this neuronal component. We categorized growth cone morphology using phalloidin staining into two different types as previously reported [28]: normal or stable growth cone, characterized by an expanding cone larger than its proximal area, containing the typical lamellipodia and filopodia structures (Fig. 2a, left); and collapsed growth cone, characterized by a thin shaped cone lacking the lamellipodial/filopodial organization (Fig. 2a, right). DIV1 hTTRWT DRG neurons presented 50% of normal growth cones, as previously seen in cultures of adult WT DRG neurons [29]. In hTTRA97S neurons we observed an approximately 27% decrease in the proportion of normal growth cones (Fig. 2b), indicating a growth cone collapse phenotype in mutant neurons.

**Fig. 2.**
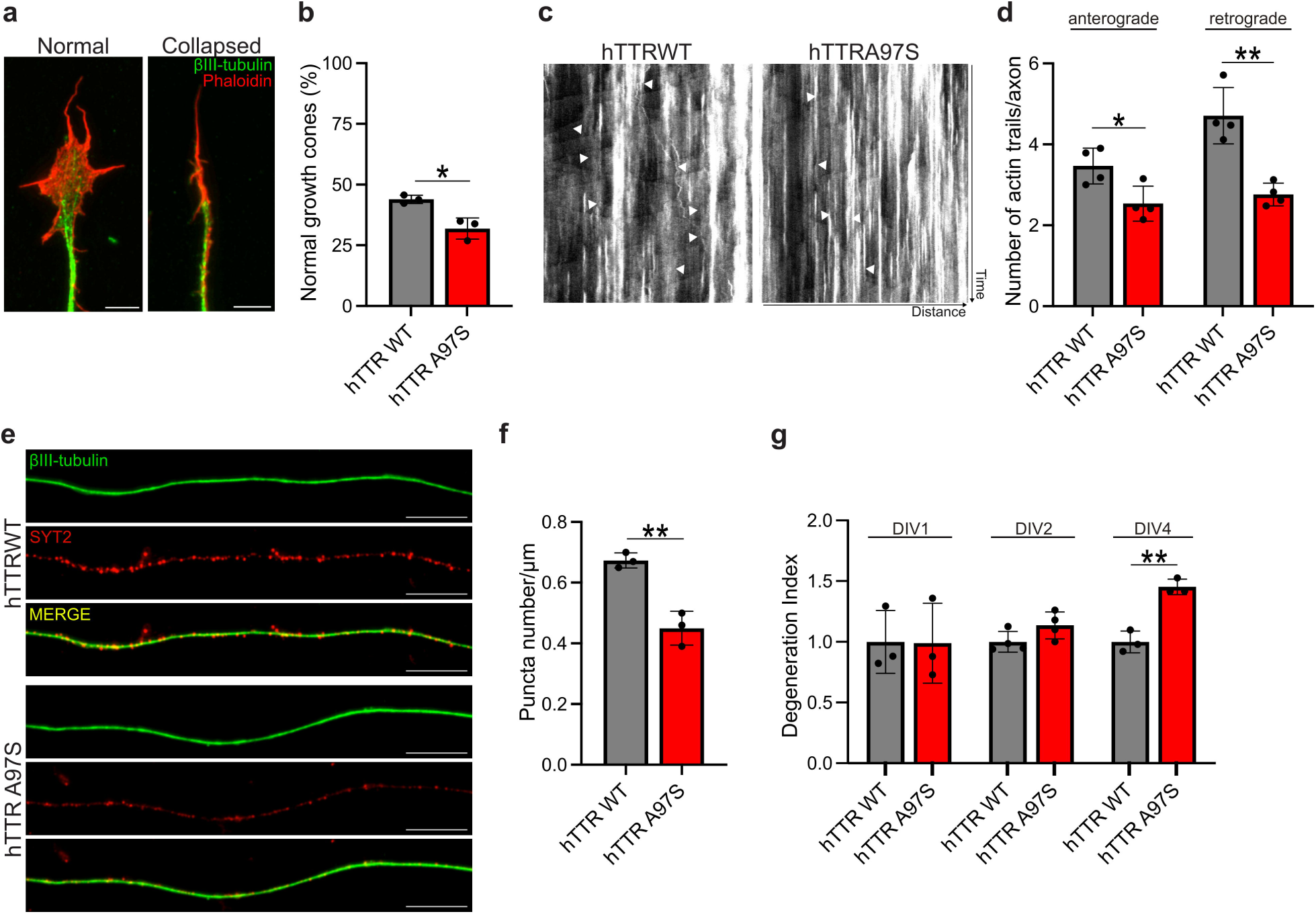
Alterations in the actin-vesicle network of hTTRA97S DRG neurons precede axonal degeneration. **a,b** Representative images of normal and collapsed growth cones of DRG neurons double-labelled with βIII-tubulin antibody (green) and phalloidin (red) **(a)** and respective quantification of the proportion of normal ones in DIV1 hTTRWT and hTTRA97S neurons **(b).** Scale bar = 5μm. Data represent mean ± SD (n=3 independent experiments). **c,d** Representative kymographs of imaged axons transfected with GFP:UTR-CH (to label F-actin) **(c)** and respective quantification of the number of retrograde and anterograde actin trails in DIV2 hTTRWT and hTTRA97S neurons **(d)**. White arrowheads mark vectorial plumes of fluorescence (actin trails). Data represent mean ± SD (n=4 independent experiments). **e,f** Representative images of DIV2 hTTRA97S and hTTRWT DRG axons double-labelled with βIII-tubulin (green) and synaptotagmin-2 (SYT2, red) **(e)** and respective quantification of the number of synaptotagmin puncta per axon length **(f)**. Scale bar = 10μm. Data represent mean ± SD (n=3 independent experiments). **g** Quantification of degeneration index in DIV1, DIV2 and DIV4 hTTRWT and hTTRA97S DRG neurons immunolabelled for βIII-tubulin. Data represent mean ± SD (n=3 independent experiments). *p<0.05, **p<0.01by Student’s t test.

Considering the axonopathy phenotype of ATTRv-PN, we analysed axonal actin organization in DIV2 hTTRA97S DRG neurons. We tracked actin dysregulation in DRG axons by evaluating actin trails, which are fast linear events of F-actin polymerization that travel throughout the axon [20]. To measure axonal actin trails dynamics we performed live imaging of DRG neurons transfected with a reporter expressing GFP fused to the calponin homology domain of utrophin (GFP:UTR-CH), which binds specifically to F-actin without significantly affecting its dynamics[30]. The nature of the actin trails displayed by control hTTRWT DRG neurons shared parameters with those previously reported in hippocampal neurons [20], such as the rate of polymerization, although the length of the trails was slightly reduced (Supplementary Table 3). Additionally, latrunculin, an F-actin depolymerizer previously shown to reduce the frequency of actin trails in hippocampal neurons [20], had a similar effect on control DRG neurons (Supplementary Fig. 2a,b). We then analyzed axonal actin trail trafficking in hTTRWT and hTTRA97S DRG neurites. While we did not find any differences regarding the length or rate of F-actin polymerization comparing the two genotypes (Supplementary Fig. 2c,d), we observed a reduction in the number of actin trails running both retrogradely and anterogradely in hTTRA97S DRG axons, supporting an axonal actin dysfunction (Fig. 2c,d).

The transport of F-actin along axons, facilitated by actin trails, was shown to support the presynaptic enrichment of F-actin, a critical process for synaptic physiology [20]. Additionally, axonal actin dynamics was related to the motion and recruitment of synaptic vesicles by regulating synaptic vesicle recycling [20, 31]. To assess whether a reduction in the number of actin trails could affect synaptic vesicles in hTTRA97S axons, we labelled recycling synaptic vesicles in DRG neurons with synaptotagmin-2, a protein embedded in the membrane of synaptic vesicles which functions as a calcium sensor that triggers the vesicle’s fusion with the presynaptic membrane. We observed a reduction in the levels of synaptotagmin-2 in hTTRA97S neurons (Fig. 2e,f), suggesting that actin defects impair synaptic physiology in hTTRA97S axons.

We followed by assessing whether the actin alterations mediated by hTTRA97S were related to axonal damage by measuring axonal degeneration in hTTRWT and hTTRA97S DRG neurons. Axonal degeneration, as determined by degeneration index measurements[32], was not different between hTTRA97S and hTTRWT neurons at DIV1, the time point of growth cone morphology analysis, or at DIV2, when axonal actin defects were observed (Fig. 2g). However, by DIV4, hTTRA97S DRG neurons exhibited axonal fragmentation, as shown by an increased degeneration index compared to hTTRWT neurons (Fig. 2g and 4c). These results demonstrate that defects in actin organization and dynamics, along with a concomitant impact on axonal synaptic physiology, precede axonal degeneration of hTTRA97S DRG neurons.

### Microtubule dynamics and axonal transport are impaired in hTTRA97S axons reflecting a generalized cytoskeleton dysfunction

Considering the crosstalk between actin and microtubules and the emerging recognition of alterations in microtubule stability and axonal transport as common features of dying-back axonopathies [33], we sought to investigate additional cytoskeleton alterations in hTTRA97S mice. Leveraging transgenic mice expressing the microtubule plus-end binding protein EB3 fused to green fluorescent protein (GFP) under the control of the neuron-specific Thy1 promoter (Thy1:EB3-GFP) [16], we examined microtubule dynamics in the sural nerve of hTTRA97S mice. We generated hTTRA97S-Thy1-EB3-GFP and hTTRWT-Thy1-EB3-GFP mice and performed *ex-vivo* live imaging of sural nerves from pre-symptomatic 9-month-old animals, an age preceding axonal degeneration. Our data revealed an increase in EB3 comet density and with no alterations in growth rate, in hTTRA97S-Thy1-EB3-GFP mice when compared to control hTTRWT-Thy1-EB3-GFP animals (Fig. 3a-c). The observed increase in EB3 comet density, combined with the lack of differences in total microtubule density assessed by electron microscopy of cross sections of hTTRWT and hTTRA97S sural nerves (Supplementary Fig. 3a), suggests an increase in dynamic microtubule plus-ends in hTTRA97S axons.

**Fig. 3.**
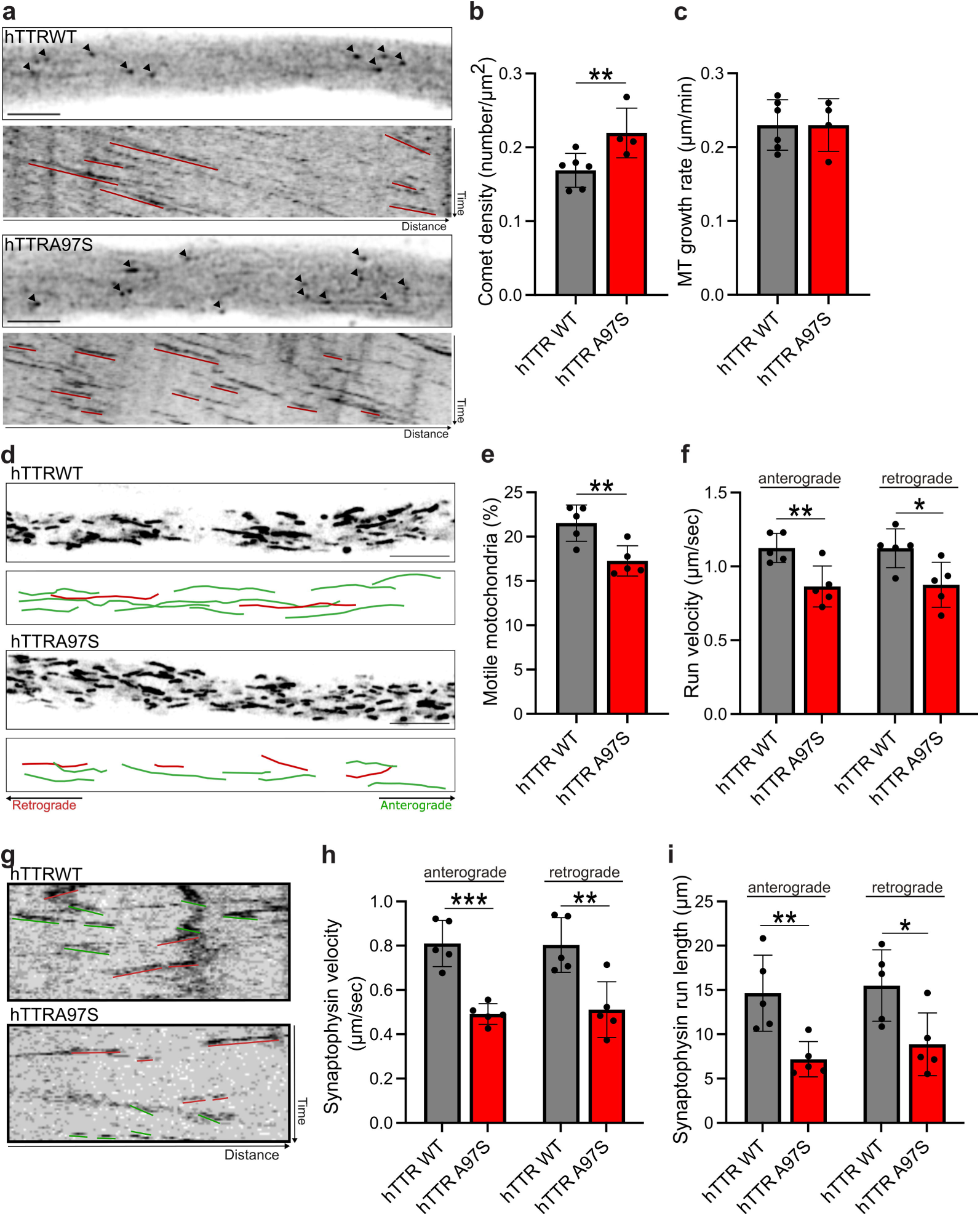
Ex vivo live imaging of hTTRA97S sural nerve axons reveals alterations in microtubule dynamicity and transport of mitochondria. **a-c** Representative EB3-GFP still images of the axonal shaft from hTTRWT-Thy1- EB3-GFP and hTTRA97S-Thy1-EB3-GFP sural nerves with correspondent kymographs **(a)**, and respective quantifications of EB3 comet density **(b)** and microtubule growth rate **(c)**. Scale bar = 5μm. Data represent mean ± SD (n=4-6 animals/genotype; 5–6 axons/animal). **d-f** Representative still images of Mito-RFP in hTTRWT-Thy1-MitoRFP and hTTRA97S-Thy1-MitoRFP sural nerve axons **(d)** and respective quantification of the percentage of motile mitochondria **(e)** and mitochondrial run velocity **(f)**. Scale bar = 10 μm. Data represent mean ± SD (n=5 animals/genotype; 5 axons/animal). **g-i** Representative kymographs of imaged neurites from DIV2 hTTRWT and hTTRA97S DRG neurons expressing synaptophysin-GFP **(g)** and respective quantification of synaptophysin velocity **(h)** and run length **(i)**. Data represent mean ± SD (n=5 independent experiments). *p<0.05, **p<0.01, ***p<0.001 by Student’s t test.

In the axonal shaft, a tight balance between microtubule stability and dynamics is critical to enable normal axon physiology. The increased microtubule dynamics in the shaft of hTTRA97S axons could therefore underlie the dysregulation of such crucial balance and potentially lead to axon pathology, particularly to a defective axonal transport. To explore this further, we analysed the transport of mitochondria by crossing hTTRA97S or hTTRWT mice with Thy1-MitoRFP mice, which express a mitochondrial sequence fused to RFP under the neuronal Thy1 promoter [34], thereby generating hTTRA97S-Thy1-MitoRFP and hTTRWT-Thy1-MitoRFP mice. Through *ex-vivo* live imaging of sural nerves from 9-month-old animals, we observed a reduction in the percentage of motile mitochondria in hTTRA97S-Thy1-MitoRFP nerves compared to the control group (Fig. 3d,e). Also, the motile mitochondria in the mutant axons exhibited reduced anterograde and retrograde velocities, as well as shorter run length in both directions (Fig. 3f; Supplementary Fig. 3b). Additionally, we assessed synaptophysin transport by transducing DRG neurons with lentivirus expressing synaptophysin-GFP. In hTTRA97S neurons, we observed a reduction in synaptophysin vesicle transport velocity, affecting both anterograde and retrograde directions, accompanied by a significant decrease in vesicle run length (Fig. 3g-i).

Together, our results suggest that hTTRA97S induces dysfunctions in both actin and microtubule cytoskeleton within peripheral axons, leading to impaired axonal transport. Importantly, these phenotypes precede axonopathy suggesting that targeting cytoskeleton damage by modulating the underlying molecular mechanisms in ATTRv-PN could potentially prevent neurodegeneration.

### Rac1 hyperactivation leads to cytoskeleton defects in hTTRA97S axons

Our previous study using an ATTRv-PN *Drosophila* model demonstrated that Rac1 silencing reverts axonal growth defects and actin growth cone alterations of mutant photoreceptor axons[12]. Building on this, we analysed the involvement of Rac1 in hTTRA97S axonal cytoskeleton alterations. Using live cell imaging with a FRET-based Rac1 biosensor [35], we detected Rac1 hyperactivation in DRG neurites shafts from hTTRA97S mice, evidenced by a 45% increase in the FRET/CFP ratio of the Rac1 FRET probe compared to hTTRWT control neurons (Fig. 4a,b). To further assess Rac1 activity, we performed pull-down assays on sciatic nerves from hTTRA97S mice. We observed an increase in Rac1 activity in hTTRA97S sciatic nerves from 9-month-old mice (Fig. 4c,d), as demonstrated by an increase in Rac1-GTP levels with no alteration in total Rac1 levels. These results demonstrate that the presence of mutant TTR in peripheral nerves results in an increase in Rac1 activity, at an age preceding disease symptomatology.

**Fig. 4.**
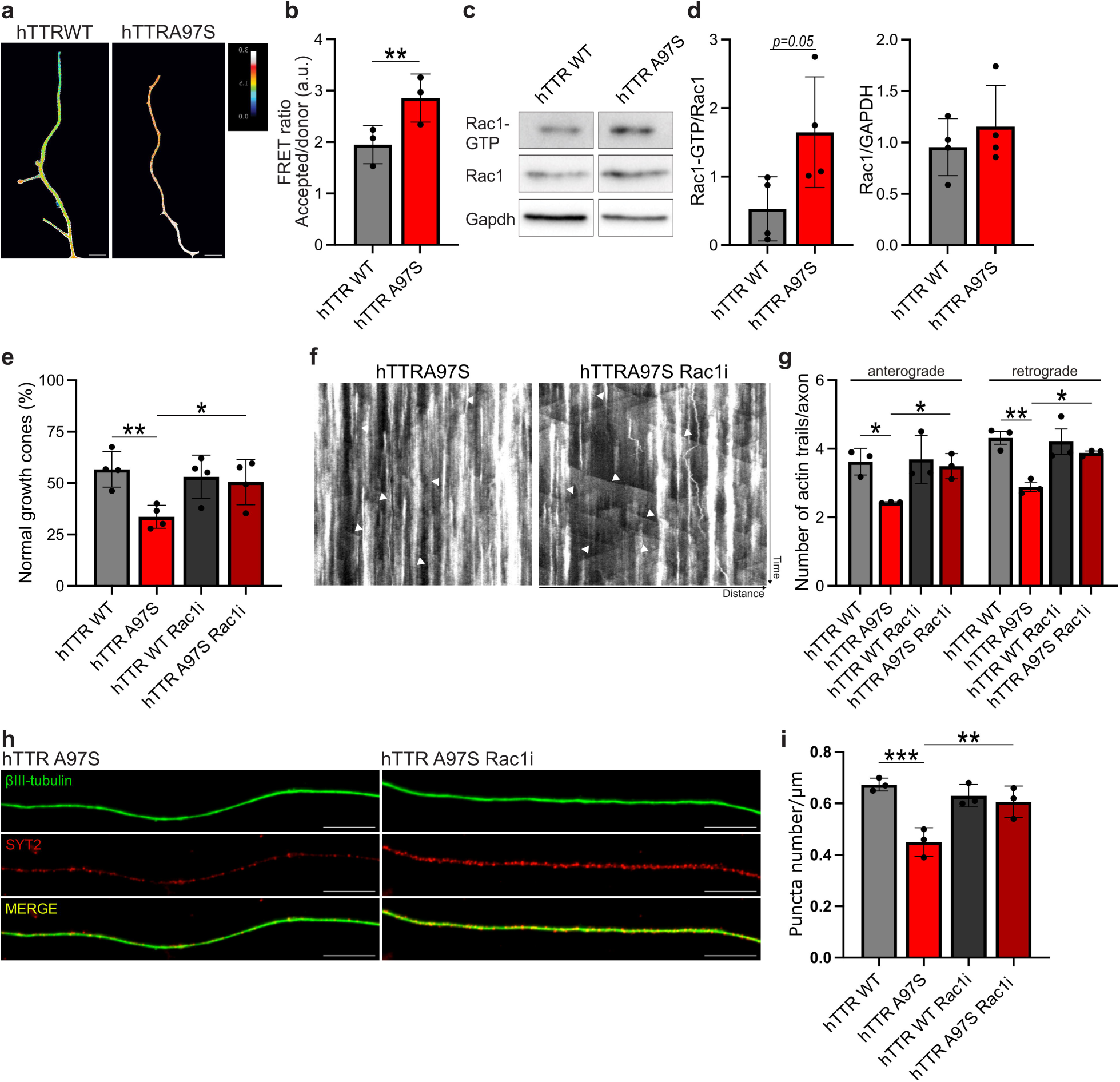
Rac1 activity is increased in hTTRA97S axons and its inhibition prevents actin-vesicle defects. **a,b** Pseudocoloured images displaying pixel value of FRET ratio in neurites from DIV2 hTTRWT and hTTRA97S DRG neurons transfected with Raichu-Rac1 biosensor **(a)**, and respective quantification **(b)**. Scale bar = 10 μm. Data represent mean ± SD (n= 3 independent experiments). **p< 0.01 by Student’s t-test. **c,d** Representative western blot analysis **(c)** and quantification **(d)** of Rac1 GTP-bound (active) levels by pull-down assay of 9- month-old hTTRWT and hTTRA97S sciatic nerves. The activated fraction was calculated by normalizing signal densities of GTP-bound to total Rac1 GTPase. Total levels of Rac1 are relative to GAPDH. Data represent mean ± SD (n= 4 animals/genotype). *p<0.05 by Student’s t-test. **e** Quantification of the percentage of normal growth cones in DIV1 hTTRWT and hTTRA97S neurons untreated or treated with 50 μM Rac1 inhibitor NSC23766 (Rac1i). Data represent mean ± SD (n= 4 independent experiments). *p<0.05, **p<0.01 by One- way ANOVA with Sidak’s multiple comparisons test. **f** Representative kymographs of imaged axons transfected with GFP:UTR-CH (to label F-actin) from DIV2 hTTRA97S DRG neurons untreated or treated with 50 μM Rac1 inhibitor NSC23766 (Rac1i). **g** Quantification of the number of retrograde and anterograde actin trails in DIV2 hTTRWT or hTTRA97S neurons untreated or treated with 50 μM Rac1 inhibitor NSC23766. Data represent mean ± SD (n= 3 independent experiments). *p<0.05, **p<0.01 by One-way ANOVA with Sidak’s multiple comparisons test. **h** Representative images of axons from DIV2 hTTRA97S DRG neurons, untreated or treated with 50 μM Rac1 inhibitor NSC23766 (Rac1i), double-labelled with βIII-tubulin (green) and synaptotagmin-2 (SYT2, red). Scale bar = 10 μm. (I) Quantification of the number of synaptotagmin puncta per axon length in DIV2 hTTRWT or hTTRA97S neurons untreated or treated with 50 μM Rac1 inhibitor NSC23766. Data represent mean ± SD (n= 3 independent experiments). **p<0.001, ***p<0.0001 by One-way ANOVA with Sidak’s multiple comparisons test.

Given the increased Rac1 activity in hTTRA97S mice and the critical role of the Rho GTPase in cytoskeleton regulation, we next examined whether Rac1 mediates actin damage in mutant neurons. We found that either silencing (Supplementary Fig. 4a-c) or blocking Rac1 activity, using the Rac1 inhibitor NSC23766 (Fig. 4e), reverted actin cytoskeleton alterations in the growth cone of hTTRA97S neurons. Additionally, we analysed whether Rac1 inhibition could similarly ameliorate the axonal actin defects observed in mutant neurons. Indeed, Rac1 inhibition restored the number of actin trails number observed in hTTRA97S DRG axons (Fig. 4f,g). The rescue of axonal actin defects was concomitant with a reversion of the decrease in synaptotagmin puncta in hTTRA97S neurons (Fig. 4h,i), reinforcing the role of Rac1 as a key regulator of actin dynamics, which subsequently impacts the presynaptic vesicle network.

We then explored whether Rac1 could also be involved in the microtubule dysregulation observed in hTTRA97S peripheral axons. For that, we evaluated microtubule dynamics in DRG neuronal cultures from hTTRA97S-Thy1-EB3-GFP and hTTRWT-Thy1-EB3-GFP mice. Mutant neurons displayed an increased EB3 comet density in axon shafts, recapitulating the data obtained *ex-vivo* in sural nerve axons (Fig. 5a,b). Implying Rac1 involvement in the alterations of microtubule dynamics, inhibition of the Rho GTPase reverted the increased EB3 comet density of hTTRA97S-Thy1-EB3-GFP DRG neurites (Fig. 5a,b). These results suggest that either Rac1 dysregulation directly affects microtubule dynamics, or that, Rac1 inhibition, by rescuing actin defects, reverts microtubule alterations. Importantly, our data pinpoints Rac1 as a critical molecular player in the cytoskeletal alterations induced by mutant TTR.

**Fig. 5.**
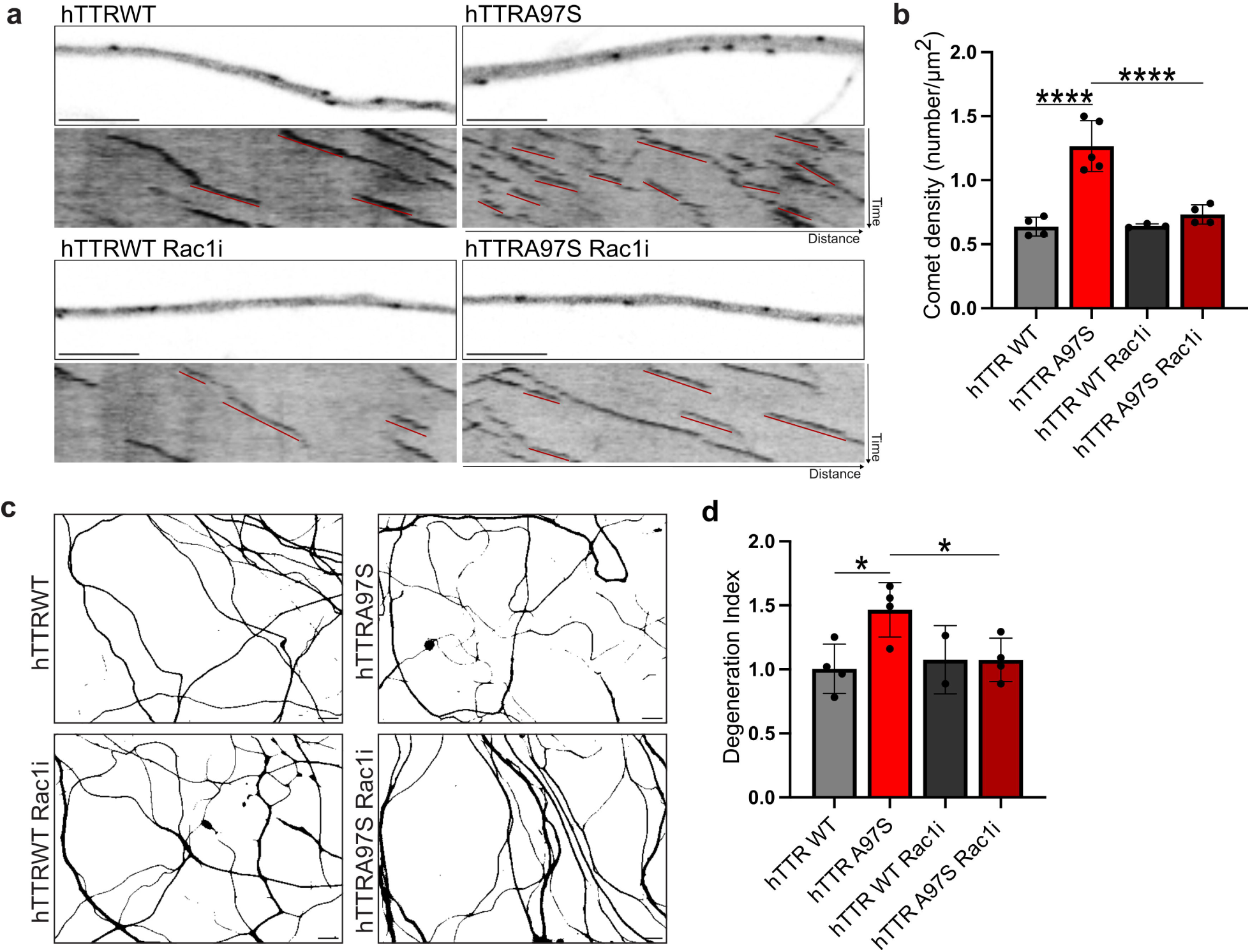
Rac1 inhibition reverts microtubule alterations and axonal degeneration in hTTRA97S neurons. **a,b** Representative EB3-GFP still images of the axonal shafts from DIV2 hTTRWT-Thy1-EB3-GFP or hTTRA97S- Thy1-EB3-GFP DRG neurons, untreated or treated with 50 μM Rac1 inhibitor NSC23766 (Rac1i) **(a)**, and correspondent quantification of EB3 comet density **(b)**. Scale bar = 5 μm. Data represent mean ± SD (n= 4 independent experiments). ****p<0.0001 by One-way ANOVA with Sidak’s multiple comparisons test. **c,d** Representative immunofluorescence images of βIII-tubulin stained DIV4 hTTRWT and hTTRA97S DRG untreated or treated with 50 μM Rac1 inhibitor NSC23766 (Rac1i) **(c)**, and respective quantification of the degeneration index **(d)**. Scale bar = 10 μm. Data represent mean ± SD (n= 4 independent experiments). *p<0.05 by One-way ANOVA with Sidak’s multiple comparisons test.

Having demonstrated that Rac1 inhibition rescues cytoskeletal damage, a phenotype that precedes axonal degeneration in mutant neurons, we evaluated the potential of inhibiting Rac1 activity to prevent amyloidogenic TTR-induced axonal degeneration. We observed that Rac1 inhibition reverted neurodegeneration in DIV4 hTTRA97S neurons, as indicated by the restoration of the degeneration index of the mutant neurons (Fig. 5c,d). These results show that blocking cytoskeleton alterations by inhibiting Rac1 in DRG neurons from an ATTRv-PN mouse model prevents axonal degeneration, identifying Rac1 as a novel therapeutic target for the disease.

### A *RACGAP1* variant is associated with disease onset in ATTRv-PN patients

The most prevalent pathogenic variant in ATTRv-PN is TTR Val30Met (ATTRV30M), and among carriers of this variant, there is significant diversity in phenotype, severity, and AO. To determine the relevance of Rac1 in human disease, we investigated whether variants in the genes encoding for Rac1 and its activators, guanine nucleotide exchange factors (GEFs), and inactivators, Rac1 GTPase-activating proteins (GAPs), might be associated with variability of AO among ATTRV30M patients. We analysed a total of 744 SNPs in RAC1*, RAC1* GAPs and *RAC1* GEFs (Supplementary Table 1). Our sample included a total of 175 patients with ATTRV30M amyloidosis belonging to 109 families, divided into two groups based on AO (Table 1). We found one SNP rs615382 in the *RACGAP1* gene associated with a late disease onset (p<0.001), specifically a mean increase in 20 years for heterozygous individuals and 34 years for homozygous individuals carrying the variant (Table 2). No SNPs in the other studied genes showed statistical significance (p>0.05) when comparing early- and late-onset individuals. Bioinformatics analysis of this SNP was performed using Ensembl (www.ensembl.org, release 110) and HaploReg (https://pubs.broadinstitute.org/mammals/haploreg/haploreg.php, HaploReg v4.2). The *RACGAP1* gene encodes RacGAP1, a specific Rac1 GAP that inhibits the Rac1 GTPase activity[14]. The variant rs615382 is located in an intronic region of *RACGAP1* and overlaps with an enhancer region.

**Table 1.**
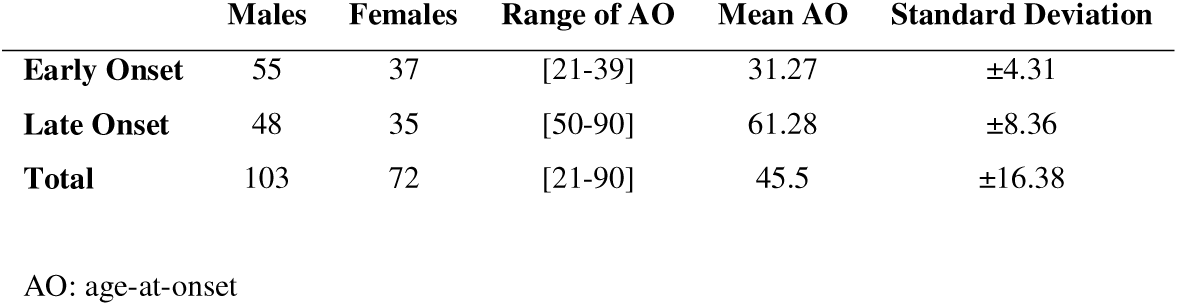
Study group demographic characteristics for Genome-Wide Array Genotyping.

**Table 2.**
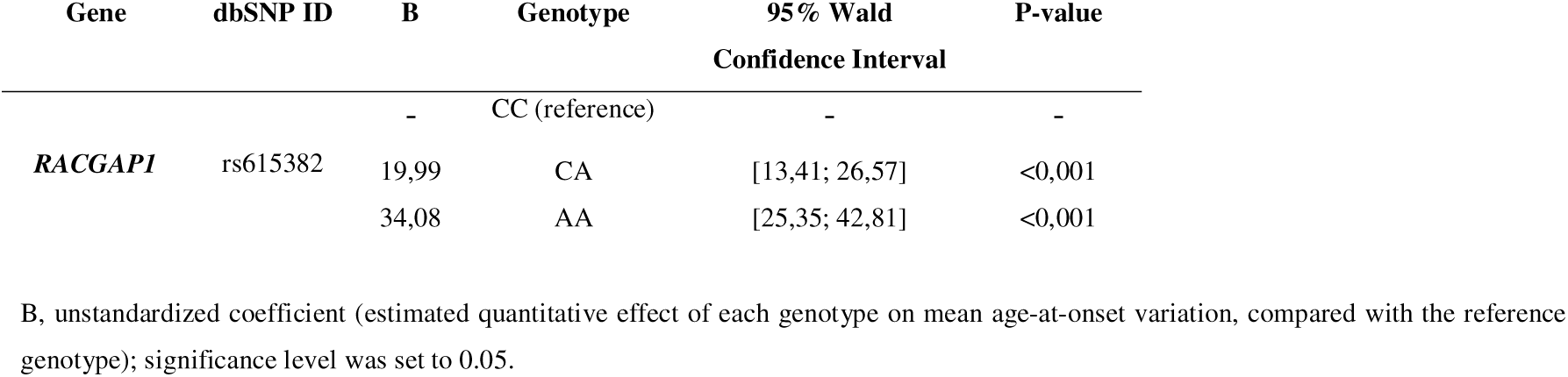
Statistical results of *RACGAP1* SNP rs615382.

Then, we analysed the influence of SNP rs615382 on *RACGAP1* expression. Plasmids encompassing the reference (C) and alternative (A) alleles of *RACGAP1* rs615382 variant were transfected into two cell lines: HEK293T, a non-neuronal cell line, and SH-SY5Y, a cell line with neuronal-like characteristics. Subsequently, the activity of the luciferase gene reporter was quantified using a luminescence assay. The luciferase activity of *RACGAP1* rs615382 A-allele showed a significant increase relative to *RACGAP1* rs615382 C-allele, with a difference of approximately 35% and 20% in SH-SY5Y and HEK293T cells, respectively (Fig. 6a,b). These results confirm that the alternative A-allele, which is associated with late-onset ATTRV30M amyloidosis, leads to higher *RACGAP1* expression. This finding provides direct evidence that rs615382 plays a critical role in modulating the expression of *RACGAP1,* suggesting that increased RacGAP1 levels are linked to the later onset of the disease. To further confirm whether increased *RACGAP1* expression is associated with late-onset ATTRV-PN, we performed qPCR in salivary gland biopsies from early and late-onset patients. Human salivary gland biopsies have been successfully used as a tool for diagnosis and research on ATTRV-PN. We detected increased *RAGGAP1* levels in late-onset patients when compared to early-onset cases (Fig. 6c), proposing that an increase in RacGAP1, which correlates with decreased Rac1 activity, is associated with a slower and less aggressive form of ATTRV-PN. Together with the data obtained with the hTTRA97S mice, these results support the neuroprotective role of Rac1 inhibition in ATTRV-PN.

**Fig. 6.**
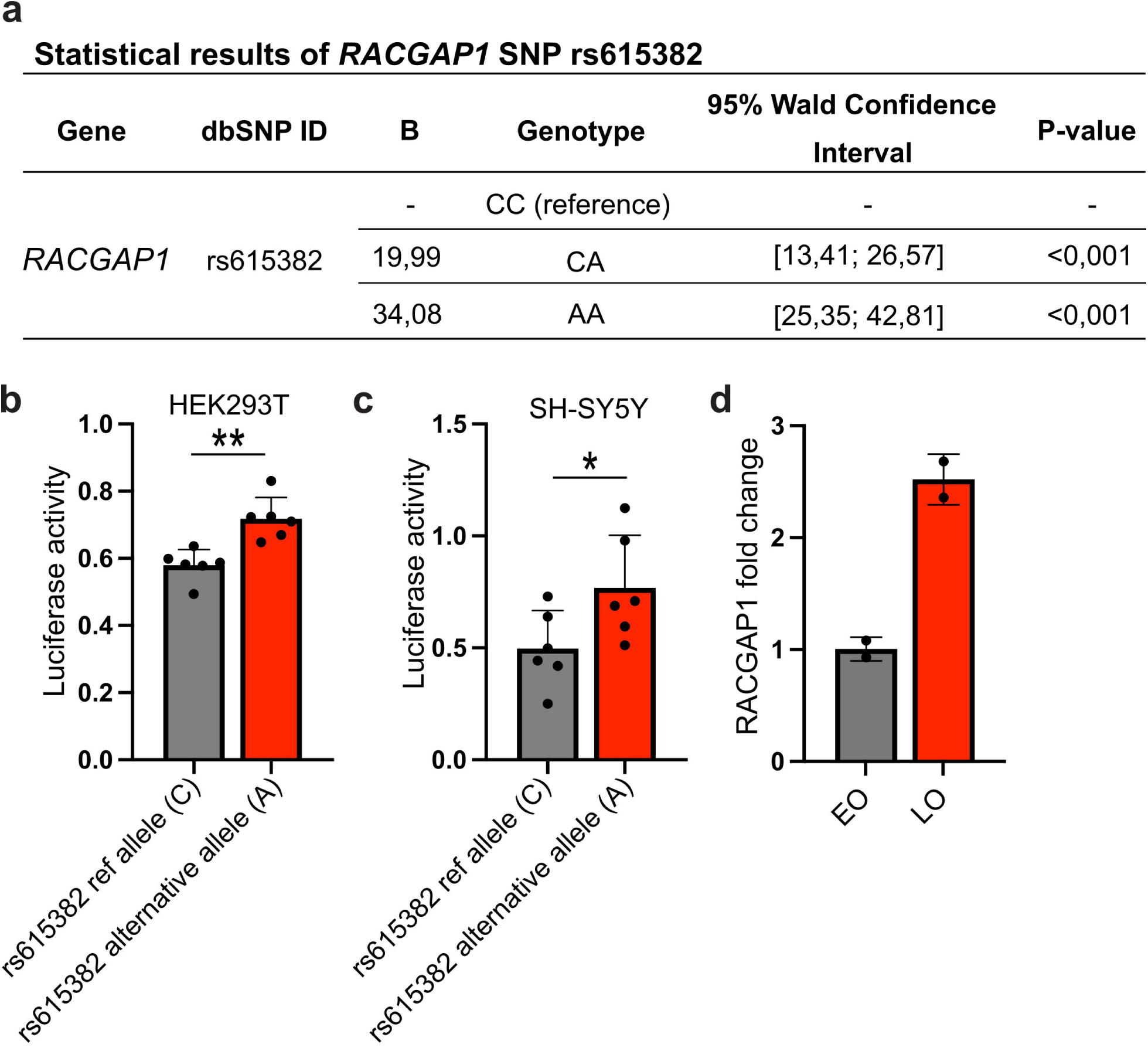
A SNP leading to the increased expression of RACGAP1, an inhibitor of Rac1 activity, is associated with late-onset ATTRV-PN **a,b** Luciferase reporter gene assays for *RACGAP1* rs615382 in HEK293T **(a)** and SH- SY5Y cells **(b)**. The activity of firefly luciferase was normalised to renilla luciferase activity and presented as a fold increase compared to the pGL3-promoter activity. Results presented as mean ± SD (n=6 independent experiments). *p<0.05, **p<0.01by Student’s t test. **c** qPCR results for *RAGGAP1* gene expression in salivary glands from early (EO) or late (LO) onset ATTRV30M patients. Data is shown as fold change in relation to EO samples. Data represent mean ± SD (n= 2 cases/disease onset).

## Discussion

The findings presented in this study demonstrate that Rac1 hyperactivation is a critical mediator of cytoskeletal dysfunction preceding axonal loss in ATTRV-PN. We hypothesize that amyloidogenic TTR, possibly through the interaction with the receptor for advanced glycation end products (RAGE), could regulate Rac1 activity. Binding of misfolded TTR to RAGE results in the activation of signaling pathways that contribute to cellular stress and inflammation [36]. Moreover, RAGE is known to trigger Rac1 signaling cascades [37], suggesting that TTR-RAGE engagement may lead to Rac1 hyperactivation. Accordingly, we observed that the RAGE-Specific Inhibitor FPS-ZM1[38] blocks TTR-induced growth cone actin alterations in hTTRA97S DRG neurons (data not shown), similarly to the Rac1 inhibitor.

Our previous work highlighted the impact of amyloidogenic TTR on the actin cytoskeleton in ATTRV30M- expressing Drosophila larvae [12]. While this model did not fully capture ATTRv-PN pathology, as patients are asymptomatic during the development and maturation of the nervous system, we now observed similar actin defects in adult hTTRA97S DRG neurons, with an increase in collapsed growth cones before axonal degeneration. Growth cone collapse is a well-known early feature of axonal degeneration, as seen in other models such as Friedreich’s ataxia, an inherited ataxia characterized by degeneration of DRG neurons [29]. We also found axonal actin defects, specifically a reduction in the number of actin trails, in hTTRA97S DRG neurons, fitting the pathologic scenario of ATTRv-PN where amyloidogenic TTR accumulates and contacts mature peripheral axons. This was a key finding, showing the role of axonal actin trails in maintaining axon health, by regulating the synaptic vesicle network [20], and the impact of their dysregulation on axonal degeneration.

In addition to unravelling actin damage as a novel player of neurotoxicity in ATTRv-PN, our results also substantiate an impairment in microtubule dynamics and axonal transport in hTTRA97S peripheral axons. Although we cannot disclose the primary defect induced by amyloidogenic TTR, our data suggests that a common pathway mediates the damage to actin and microtubules in ATTRv-PN, driven by Rac1. Mechanistically an increase in Rac1 activity might contribute to the cytoskeleton defects observed in hTTRA97S axons. Rac1 is a well-established regulator of actin polymerization and branching, facilitating lamellipodia and filopodia formation through downstream effectors such as the WAVE complex and the Arp2/3 actin-nucleating complex [39]. In the context of ATTRv-PN, Rac1 overactivity could lead to aberrant actin branching and disorganized filament networks, contributing to the growth cone collapse phenotype. Interestingly, in our proteomic analysis, we found Arpc1a, one of seven subunits of the human Arp2/3 protein complex as being upregulated in hTTRA97S axons. Additionally, despite Rac1 being primarily recognized as an Arp2/3 activator, the Rho GTPase was shown to be able to both activate and inhibit Arp2/3-driven actin branching and polymerization [40], which would also impact growth cone morphology. Our results also show that the reduction of actin trails in hTTRA97S DRG axons is mediated by the overactivity of Rac1. Actin trails were shown to be formin-dependent and Arp2/3-independent in hippocampal neurons [20]. While we did not detect alterations in formins in our proteomic analysis, it is possible that an increase in Arp2/3 resulting in a limited G-actin pool would inhibit formins, as few actin monomers would be available for polymerization. Additionally, Rac1 can regulate microtubule dynamics through pathways like PAK1 activation, which inactivates stathmin via phosphorylation, promoting microtubule polymerization [41]. Future work will be required to detail the molecular mechanism of Rac1 involvement in ATTRV-PN.

Importantly, we validated the relevance of Rac1 in human ATTRV-PN, since the variant rs615382 in the *RACGAP1 gene* was significantly associated with delayed disease-onset in ATTRV30M patients. The SNP identified was suggested to increase *RACGAP1* expression by disrupting the Recombination Signal Binding Protein for Immunoglobulin kappa J region (RBPJ) binding site to gene corepressors, such as Silencing- Mediator for Retinoid/Thyroid hormone receptor (SMRT) and Msx2-interacting nuclear target (MINT) [42–45], what would prevent the protein from repressing transcription [46]. Results from our reporter gene assays and qPCR data with salivary glands from ATTRV-PN patients, substantiate this hypothesis, demonstrating that the alternative allele (A) of rs615382 indeed elevates *RACGAP1* expression when compared to the reference allele (C). RacGAP1 has been studied primarily for its role in cell division and cytoskeletal dynamics and our work underscores the importance of RacGAP1 in modulating neurodegenerative processes. Studies in zebrafish with a loss of function *RACGAP1* mutation have demonstrated that RacGAP1 is required for the long-term survival of motor neurons and the cytoskeletal organization of sensory axons [47]. Also, the fact that we found the rs615382 SNP, and increased *RAGGAP1* expression associated with a later onset of ATTRV30M, suggests that an increase in RacGAP1 might be translated to Rac1 inactivation leading to a lower rate of neurodegeneration. Rac1 plays essential roles in many tissues, including the cardiovascular and immune systems, making targeted therapeutic interventions difficult without off-target effects. Although Rac1 inhibitors like NSC23766 have shown promise in our experimental model, their clinical use is challenging. Clinical trials of Rac1 inhibitors are still in the early stages, and achieving therapeutic specificity remains a significant hurdle. Nevertheless, our identification of the *RACGAP1* variant in ATTRv-PN patients, which enhances the expression of this Rac1 inactivator, offers a potential strategy to modulate Rac1 activity more selectively. We are currently working on AAV [48] and nanoparticle [49] strategies targeting the PNS to modulate *RACGAP1 in vivo*. This approach represents significant progress from current treatments, which predominantly focus on TTR stabilization or gene silencing, and do not address the downstream effects of TTR deposition on neuronal health.

In summary, we show that Rac1 inhibition preserves cytoskeletal integrity and prevents axonal degeneration in a mouse model for ATTRv-PN. Importantly, we establish that a variant in the gene encoding the Rac1-specific inactivator RacGAP1, which leads to its increased expression, is associated with a late-onset form of the disease. This work supports the neuroprotective effect of Rac1 inactivation in ATTRv-PN, offering new insights into the disease’s molecular mechanism and identifying promising targets for therapeutic intervention. Future research will be required to translate these findings into pre-clinical studies, with the ultimate goal of improving outcomes for patients with ATTRv-PN.

## Methods

### Subjects and Study Design

DNA samples from 175 ATTRv-PN ATTRV30M amyloidosis patients were collected at Corino de Andrade at Centro Hospitalar Universitário de Santo António (UCA-CHUdSA) and stored at Centro de Genética Preditiva e Preventiva (CGPP-IBMC, Porto) biobank, as authorized by the CNPD (National Commission for Data Protection). Salivary gland biopsies from ATTRV30M patients were also collected at UCA-CHUdSA. All patient samples were clinically characterized and patients were observed by the same group of neurologists. Age-at-onset (AO) of each patient was also registered. According to the literature, the samples were divided according to AO into early-onset (EO), when symptoms appeared before the age of 50 and late-onset (LO) if symptoms appeared only after that age [15]. The study was conducted on 92 DNA samples of EO patients, 83 DNA samples of LO patients, 2 salivary glands samples of EO patients and 2 salivary glands of LO patients. The Ethics Committee of CHUdSA approved the study and all participants gave their written informed consent.

### Animals

Mice were handled according to European Union and National rules. All animals were maintained under a 12h light/dark cycle and fed with regular rodent chow and water *ad libitum*. Genotypes were determined from ear- extracted genomic DNA [13]. Human TTR knock-in mice, carrying either wild-type human TTR or A97S human TTR were generated by replacing the m*Ttr* gene with the h*TTR* gene without altering the promoter and enhancer sequences of m*Ttr*, as previously reported [13]. Mice were maintained in a C57BL/6 background and heterozygotes with the genotypes hTTRwt/mTtrwt (abbreviated as hTTRWT; controls) and hTTRA97S/mTtrwt (abbreviated as hTTRA97S) were used in this study. For measuring microtubule dynamics and axonal transport, homozygous hTTRA97S/hTTRA97S and hTTRWT/hTTRWT mice were crossed with either Thy1:EB3-GFP [16] or Thy1-MitoRFP mice [17] (kindly provided by Dr Thomas Misgeld, Technical University of Munich, Germany), generating hTTRA97S-Thy1-EB3-GFP and hTTRWT-Thy1-EB3-GFP mice or hTTRA97S-Thy1- MitoRFP and hTTRWT-Thy1-MitoRFP mice, respectively.

### Myelinated axons quantification

Sural nerves from 9-month-old hTTRWT and hTTRA97S were collected and fixed overnight at 4 °C with a solution of 2.5% glutaraldehyde and 2% paraformaldehyde (PFA) in 0.1 M sodium cacodylate buffer (pH 7.4), and post-fixed with 2% osmium tetroxide in 0.1 M sodium cacodylate buffer for 2 h at RT and stained with 1% uranyl acetate for 30 min. Samples were dehydrated and embedded in Epon (Electron Microscopy Sciences). Cross sections were cut at 500 nm using a diamond knife in an RMC Ultramicrotome (PowerTome, USA) and contrasted with toluidine blue. Image acquisition was performed with a brightfield microscope Leica DM2000 LED (Leica Microsystems, Germany) using a 40x objective. The total number of myelinated axons was determined in each cross section, using the Cell Counter plugin from ImageJ, and normalized by the total nerve area.

### Quantification of skin innervation

Mice were perfused with 2% Periodate-Lysine-Paraformaldehyde (PLP). The footpad tissues were postfixed with PLP for 2h and cryosectioned into 30μm free-floating sections perpendicular to the dermis. For immunohistochemistry, tissue sections were permeabilized with 0.3% Triton X-100 for 10 min, blocked with 10% BSA in 0.3% Triton in PBS for 1 h at RT, and incubated with rabbit anti-PGP9.5 (1:1,000; Abcam, ab108986) in blocking buffer overnight at 4°C. Incubation with anti-rabbit Alexa Fluor-conjugated secondary antibody was performed for 1 h at RT. Epidermal innervation was quantified following established protocols, and the slides were coded to ensure that measurements were blinded [13]. Any nerve fiber ascending from the subepidermal plexus with branches inside the epidermis was counted as one, while branches in the dermis were counted as separate fibers. ImageJ was used to measure the total length of the epidermis along the upper margin of the stratum corneum. Intraepidermal nerve fiber density was derived as the number of fibers per millimeter of epidermal length.

### von Frey test

For von Frey hair testing, animals were acclimatized for 20 min in a chamber with a wire-mesh bottom allowing access to hind paws. For the test, retractable monofilaments (Aesthesio, Precise Tactile Sensory Evaluator, 37450-275) were used to apply a force to the mid-plantar surface of hind paws. Paw withdrawal or abrupt movement were considered positive responses. The withdrawal threshold equaled the weakest force to elicit paw withdrawal in 50% or more of the trials (n = 5 trials). The force of withdrawal is presented as the average value of the right and left hind paws.

### TTR western blot analysis

9-month-old hTTRWT and hTTRA97S mice were perfused with PBS and sural nerves collected. Protein lysates were prepared in ice-cold RIPA lysis buffer (Merck, R0278), supplemented with a cocktail of protease inhibitors (Protease Inhibitor Mix, Merck, GE80-6501-23) and 1 mM sodium orthovanadate, and then sonicated in a water bath sonicator (Bioruptor Plus, Diagenode, Belgium). 5 μg of protein extracts were separated under denaturing conditions in a 12% agarose gel, transferred to Amersham Protran Premium 0.45 μm nitrocellulose membranes (GE Healthcare Life Sciences), and blocked in 5% non-fat dried milk in TBS-T for 1 h at RT. Membranes were probed overnight at 4°C with primary antibodies: rabbit anti-human TTR (1:1,000; DAKO, A002); custom-made rabbit anti-mouse TTR antibody (1:500; produced against recombinant mouse TTR); and rabbit anti-mouse vinculin (1:3,000; Thermo Fisher, 42H89L44). Subsequently, incubation with the secondary antibody anti-rabbit IgG-HRP (1:10,000; Jackson Research, 111-035-003) was performed for 1 h at RT. Immunodetection was achieved by chemiluminescence using ECL (Millipore, WBLUR0500) and quantified using Image Lab software (version 6.0.1).

### Proteomics

Proteomics was performed by the i3S Proteomics facility. For preparation of total protein extracts, sural nerves from 9-month-old hTTRWT and hTTRA97S were lysed in 100 μL ice-cold RIPA lysis buffer (TE buffer pH 8.0 (10 mM Tris and 1 mM EDTA), 140 mM NaCl (Acros Organics, 207790010), 0.1% SDS (stock 10%, NZYTech, MB11601), 1% Triton X-100 (Merck, T9284), supplemented with a cocktail of protease and phosphatase inhibitors (Halt^TM^ Protease & Phosphatase Inhibitor Cocktail, Thermo Scientific^TM^, 78440). Samples were processed using the solid-phase-enhanced sample-preparation (SP3) method and enzymatically digested with Trypsin/LysC (2µg) overnight at 37 °C with agitation at 1,000 rpm. Peptide concentration within each sample was determined through fluorescence measurements. For protein identification and label-free quantification, a nanoLC-MS/MS setup was employed, which consisted of an Ultimate 3000 liquid chromatography system coupled with a Q-Exactive Hybrid Quadrupole-Orbitrap mass spectrometer (Thermo Scientific, Germany). The UniProt Mus Musculus Proteome Database (2022_05 version, 17132 entries) was used as a reference for protein identification. Proteomics data was analyzed using Proteome Discoverer software (version 3.0.1.27, Thermo Scientific, USA). To elucidate the distinct protein levels in control and hTTRA97S mutation groups, a series of data processing steps were applied to the raw data, including a minimum requirement of 2 unique or razor peptides, defining upregulated proteins based on abundance ratios (hTTRA97S/hTTRWT) of ≥1.50 and downregulated proteins with ratios of ≤0.67, setting a significance threshold for the abundance ratio p-value at ≤0.05 and determining that the protein needed to be detected in at least 2 samples within each experimental group for inclusion in the analysis. Protein functional enrichment analysis of the altered proteins in hTTRA97S versus hTTRWT sural nerves was performed with PANTHER 19.0 (http://pantherdb.org/). The online software tool ClustVis (https://biit.cs.ut.ee/clustvis/) was used to generate the heat map. For this purpose, protein abundance ratio values were transformed using log2.

### DRG primary cultures, plasmid expression and drug treatment

Primary cultures of DRG neurons from 4- to 8-weeks-old mice (hTTRA97S and hTTRWT for all experiments except for in vitro EB3 dynamics in which hTTRA97S-Thy1-EB3-GFP and hTTRWT-Thy1-EB3-GFP were used) were performed as previously described [18, 19]. DRG neurons were plated in 20 μg/mL PLL + 5 μg/mL Laminin coated glass surfaces in DMEM/F12 (Merck, D8437) supplemented with B27 (Gibco), 1% penicillin/streptomycin (Gibco), 2 mM L-glutamine (Gibco), and 50 ng/mL NGF (Millipore, 01-125)- DRG medium, at 37°C and 5% CO_2_.

For growth cone morphology assessment and synaptotagmin quantification, neurons were plated at a density of 5,000 cells in 24-well plates and cultured for 24 h (DIV1) and 48 h (DIV2), respectively. For live imaging of actin trails, DRG neurons were transfected, using the 4D Nucleofector Amaxa system (Lonza, Barcelona, Spain), with 0.5 μg of pIRESneo3+GFP:UTR-CH (kindly provided by Dr Jorge Ferreira, i3S), expressing the calponin homology (CH) domain of utrophin (UTR) which binds specifically to F-actin [20], plated at a density of 20,000 cells in coated 35 mm glass bottom μ-dishes (iBidi), and cultured for 48h. For live imaging of synaptophysin transport DRG neurons, platted at a density of 15,000 cells in coated 35 mm glass bottom μ- dishes (iBidi), were transduced at DIV1 with the lentivirus f(syn)-pSyp-GFP-w (produced at the Viral Core Facility Charité – Universitaetsmedizin Berlin, BLV-177) [21] and imaged at DIV2. For EB3 dynamics, hTTRA97S-Thy1-EB3-GFP and hTTRWT-Thy1-EB3-GFP neurons were plated a density of 15,000 cells in coated 35 mm glass bottom μ-dishes (iBidi) and imaged at DIV2.

For FRET experiments, DRG neurons were transfected using the 4D Nucleofector Amaxa system with 1 μg of Raichu Rac1 plasmid [22]. Cells were left in suspension for 24 h and were subsequently plated in a coated 4- well ibidi slide (iBidi) and cultured for an additional 24 h.

For experiments with Rac1 inhibition, cells were plated in the presence of 50 μM NSC23766 (Tocris Bioscience). For transfection-mediated Rac1 silencing, DRG neurons were nucleofected with 0.75 μg of the shRNA Rac1 (TRCN00000551888, Sigma-Aldrich) or shRNA scramble (Sigma-Aldrich) and 1.125 μg of EGFP-C1 plasmid. After transfection, cells were left in suspension for 24 h and then plated in coated 24-well plates at a density of 5,000 cells and cultured for 24 h.

For degeneration index assessment, DRG neurons were plated in coated 24-well plates at a density of 10,000 cells in DRG medium and: i) DIV1 analysis: fixed 24 h after plating; ii) DIV2 analysis: plated with 60 μM 5- Fluoro-2’- deoxyuridine (FluoU) and fixed 48 h later; iii) DIV4 analysis: plated with 60 μM FluoU, refreshed with 60 μM FluoU (and 50 μM NSC23766, when testing Rac1 inhibition) at DIV2 and fixed at DIV4.

### Validation of Rac1 silencing

N1E-115 cells (Sigma-Aldrich), an adrenergic cell line derived from the mouse neuroblastoma C1300 tumor, were grown in DMEM supplemented with 2mM L-glutamine, 10% FBS and 1% P/S in uncoated 24 well plates at a density of 150,000 cells/well. At DIV1, cells were transfected with 0.5 μg of the shRNA Rac1 or shRNA scramble (Sigma-Aldrich) using lipofectamine 2000. At DIV3, protein extracts were prepared in lysis buffer (0.3% Triton X-100, 1x protease inhibitor Cocktail and 1mM Sodium orthovanadate). 20 μg of protein extracts were separated under denaturing conditions in a 12% agarose gel, transferred to Amersham Protran Premium 0.45 μm nitrocellulose membranes (GE Healthcare Life Sciences), and blocked in 5% non-fat dried milk in TBS-T for 1 h at RT. Membranes were probed overnight at 4°C with primary antibody mouse anti-Rac1 (1:2,000; Abcam, ab33186) and subsequently with the secondary antibody anti-mouse IgG-HRP (1:10,000; Jackson Research, 115-035-003) in TBS-T for 1 h at RT. Immunodetection was performed by chemiluminescence using ECL (Millipore, WBLUR0500) and quantified using ImageJ software.

### Live imaging of DRG neurons

Live imaging of actin trails on GFP:UTR-CH transfected neurons was performed in a Nikon Eclipse Ti (Nikon, Japan) inverted epifluorescence microscope equipped with an IRIS 9 camera (Teledyne Photometrics, USA) with the following settings: PL APO LAMBDA 60x/1,4 Oil DIC WD 0.13 mm objective, 12% LED power and ND4 filter. Transfected neurites were identified based on morphology and only neurons with unambiguous morphology were selected for imaging. GFP:UTR-CH was typically imaged at 1 frame/300 sec for 2 min. To analyze GFP:UTR-CH kinetics, kymographs were performed using the Fiji KymoResliceWide plugin (distance- x axis; time-y axis). Actin trails appear in these kymographs as faint fluorescent lines, referred to as “fluorescent plumes” [20]. Starting and end positions of the traces and number of trails were defined using the Fiji Cell Counter.

Imaging of synaptophysin vesicles was conducted using a confocal Leica SP8 microscope (Leica Microsystems) and Leica Application Suite X (LAS X) software with a PL APO 63x / 1.30 Glycerol objective. Image acquisition was performed over a 2 min period with frames captured at 2 sec intervals. For quantification, the MTrackJ plugin from ImageJ was used and all parameters were assessed by manually tracking motile vesicles during 60 frames of each video.

For FRET, imaging was performed using a Leica DMI6000B inverted microscope. High-speed low vibration external filter combinations (CFP excitation plus CFP emission [CFP channel], and CFP excitation plus YFP emission [FRET channel]) were mounted on the microscope (Fast Filter Wheels, Leica Microsystems). A 440- 520nm dichroic mirror (CG1, Leica Microsystems) and an HCX PL APO 63X 1.3NA glycerol immersion objective were used for CFP and FRET images. Images were acquired with 2Å∼2 binning using a digital CMOS camera (ORCA-Flash4.0 V2, Hamamatsu Photonics). Shading illumination was online corrected for CFP and FRET channels using a shading correction routine implemented for the LAS AF software. CFP and FRET images were sequentially acquired using different filter combinations (CFP excitation plus CFP emission (CFP channel), and CFP excitation plus YFP emission (FRET channel), respectively). Ratiometric FRET was calculated as acceptor/donor, as described in [23]. Image analysis was performed semi-automatically using an in-house developed macro for Fiji that allows for batch-processing of files (code available at https://github.com/mafsousa/2DFRETratiometrics). The main workflow consists of the following steps: 1) a preprocessing stage including shading correction and background subtraction (either by using a background image or by subtracting user-defined background mean intensity values); 2) cell segmentation using a user-selected channel, the best threshold algorithm and an option for user-dependent refinement; and 3) ratiometric analysis by dividing selected preprocessed channels in the segmented cell with final ratio images represented with a Royal LUT.

### Ex-vivo live imaging of Microtubule Dynamics and Axonal Transport

To analyze microtubule dynamics and axonal transport in *ex vivo* live imaging, sural nerves were collected from hTTR (A97S or WT)-Thy1-EB3-GFP and hTTR (A97S or WT)-Thy1-MitoRFP mice, respectively, and placed in a 35 mm µ-Dish (iBidi) in pre-heated phenol-free Neurobasal medium (Thermo Fisher Scientific). To evaluate EB3 dynamics in vitro DIV2 hTTRA97S-Thy1-EB3-GFP and hTTRWT-Thy1-EB3-GFPDRG neurons were imaged. Imaging of EB3 comets and mitochondria movement was conducted using a Leica SP8 microscope (Leica Microsystems) and Leica Application Suite X (LAS X) software with a PL APO 63x / 1.30 Glycerol objective. Image acquisition was performed for 2 min with frames captured at each 2 sec. For the quantification of EB3 dynamics, kymographs were generated using the Fiji KymoResliceWide plugin (distance: x-axis; time: y-axis). The start and end positions of the kymograph slopes and the number of comets were defined using the Cell Counter plugin. Mitochondrial axonal transport was analyzed using the MTrackJ plugin from ImageJ and all parameters were assessed by manually tracking mitochondria during 30 frames of each video. To be considered, mitochondria needed to be tracked in at least 5 consecutive frames. The percentage of motile mitochondria was determined by tracking both motile and immobile mitochondria.

### Immunocytochemistry

DRG neurons were fixed with cytoskeleton preservation fixative, PHEM (4% PFA, 4% sucrose, 0.25% Glutaraldehyde, 0.1% Triton X-100, 300 mM PIPES, 125 mM HEPES, 50 mM EGTA and 10 mM Magnesium Chloride), permeabilized with 0.2% Triton X-100 for 5 min, quenched with 200 mM Ammonium Chloride for 5 min and blocked with 2% Fetal Bovine Serum (FBS), 2% BSA and 0.2% Fish Gelatine in PBS for 1 h at RT.

Incubation of primary antibodies was performed in 10% blocking buffer overnight at 4°C. The following antibodies were used: mouse anti–βIII-tubulin (1:2,000; Promega, G7121) and rabbit anti-synaptotagmin-2 (1:500; SYSY, 105 222). Alexa Fluor-conjugated secondary antibodies were incubated for 1 hat RT in 10% blocking buffer. Actin was labelled with the probe-conjugated dye Rhodamine-conjugated Phalloidin (1:100; Life Technologies, R415), in parallel with the secondary antibody.

### Imaging and quantification

Immunocytochemistry images were acquired using an epifluorescence microscope Zeiss Axio Imager Z1 microscope with an Axiocam MR3.0 camera and Axiovision 4.7 software, using a Plan-Apo 63X 1.4A objective (growth cone analysis), an EC-Plan-Neofluar 40x/1.30 Oil Ph3 objective (synaptotagmin imaging), or an EC Plan Neofluar 20X 0.50NA objective (degeneration index).

Growth cones from DRG neurons labelled with βIII-tubulin and phalloidin were qualitatively categorized according to their actin organization: the normal pattern containing the typical lamellipodia and filopodia structures, organized in a star-shaped morphology, and the collapsed pattern characterized especially by the absence of lamellipodia as well as growth cones with dystrophic actin morphologies, such as actin patches. Approximately, 100 growth cones were analysed in each condition using ImageJ software.

For synaptotagmin quantification, DRG neurites were randomly selected, and subsequent analysis was performed using Fiji software. A threshold was applied to the image and adjusted according to the fluorescent signal, allowing synaptotagmin particle analysis. Puncta number per neurite length was quantified, by identifying puncta along at least 40 μm of neurite per image.

To assess axonal fragmentation through degeneration index quantification, the area occupied by the axons (total axonal area) and degenerating axons (fragmented axonal area) was analysed using the particle analyser algorithm of ImageJ (size of small fragments = 20–10,000 pixels). The degeneration index was calculated as the ratio between fragmented and total axonal areas.

### Rac1 pull down

Dissected sciatic nerves from hTTRWT and hTTRA97S mice were homogenized in lysis buffer (10 % glycerol, 50LJmM Tris-HCl pH 7.4, 100 mM NaCl, 1 % NP-40, 2 mM MgCl2 and protease inhibitor cocktail [Sigma]). 10 % of lysate volume was reserved to determine total protein amounts, and the remaining protein mixture was immunoprecipitated with the respective bait substrate (GST-tagged p21-activated kinase-binding domain [GST-PAK-PBD]), that was immobilized on Glutathione Sepharose beads (GE Healthcare). Bait-couple beads were incubated with lysates incubated with overnight at 4 °C, washed with 10 packed volumes of lysis buffer, and bound proteins were eluted in GLB buffer (150 mM Trizma Base, 6% SDS, 0.05% Bromophenol Blue, 30% glycerol and 6 nM EDTA pH 8.8) with incubation at 95°C for 10 min. Samples were resolved on a 12% SDS PAGE gel followed by standard western blot and Ponceau *S* staining to confirm uniform pull-downs before detection with the relevant antibodies: mouse anti-Rac1 (1:500; Abcam, ab33186) and mouse anti-GAPDH (1:100,000; HyTest, #6C5).

### DNA Extraction and Genome-Wide Array Genotyping in ATTRv-PN patient samples

Genomic DNA extraction from peripheral blood samples was performed using QIAamp® DNA Blood Mini Kit [24]. DNA quantification was performed using Nanodrop One. The genotyping was attained with the Axiom™ Precision Medicine Diversity Array (PMDA, Affymetrix) and the GeneTitan Multi-Channel (MC) Instrument (Thermo Fisher Scientific, Waltham, MA, USA). Genotyping raw data was analysed with the Axiom^TM^ Analysis Suite version 5.1 (Applied Biosystems), using the Best Practices Workflow with default settings.

Data referring to single nucleotide polymorphisms (SNPs) of the *RAC1* gene or its guanine nucleotide exchange factors (GEFs) and GTPase-activating proteins (GAPs) were exported and analysed. The known RAC1-specific GAPs and GEFs are listed in Supplementary Table 1.

### Plasmid cloning of RACGAP1 variants

Plasmids were generated by inserting the genomic sequences (1186 base pairs of length) surrounding the *RACGAP1* rs615382 variant (NC_000012.11:g.50412911 C>A) into the pGL3-promoter vector (Promega, Fitchburg, WI, USA). This region of *RACGAP1* was amplified via PCR from the genomic DNA of a LO patient harboring the variant (rs615382 alternative allele A). The PCR product was then purified using the Zymoclean Gel DNA Recovery Kit (Zymo Research, Irvine, CA, USA) and subsequently cloned into the pGL3-promoter vector downstream of the firefly luciferase reporter gene by Gibson Assembly (New England Biolabs, Ipswich, MA, USA).

To obtain the reference allele configuration (rs615382 reference allele C), site-directed mutagenesis was performed using the Q5 Site-Directed Mutagenesis Kit (New England Biolabs, Ipswich, MA, USA), adhering to the protocol provided by the manufacturer, and the following primer pairs were used: forward primer 5′- CTCCCCTTCCcACAGCATAATCACTAAACC-3′ and reverse primer 5′-GAACCAGAGGTGATTC-3′.

Constructs sequence was confirmed by Sanger sequencing.

### Dual-luciferase reporter gene assays

SH-SY5Y cells (DSMZ, Braunschweig, Germany) were cultured in a DMEM/F-12 high glucose GlutaMAX™, supplemented with 10% fetal bovine serum (FBS) and 1% antibiotic-antimycotic solution (Gibco, Thermo Fisher Scientific, Waltham, MA, USA). HEK293T cells (ATCC) were cultured in DMEM high glucose GlutaMAX™ medium, also supplemented with 10% FBS and 1% antibiotic-antimycotic (Gibco, Thermo Fisher Scientific, Waltham, MA, USA). Dual-luciferase reporter gene assays were performed as described in [25]. Briefly, for a period of 48 h, HEK293T and SH-SY5Y cells underwent transient transfection with vectorspGL3- promoter-RACGAP1-rs615382(C), pGL3-promoter-RACGAP1-rs615382(A), pGL3-control, or pGL3- promoter (150 ng per well) in 96-well white plates (CELLSTAR® plates with µClear® bottom; Greiner Bio-One, Kremsmünster, Austria). To monitor transfection efficiency, 15 ng of the internal control pRL-CMV renilla vector (Promega, Fitchburg, WI, USA) was co-transfected into cells. Transfection with the reagent DreamFect Gold (OZ Biosciences, Marseille, Provence-Alpes-Cote d’Azur, France) was performed following the manufacturer’s guidelines. Complexes were added to each well with 100 µL of medium devoid of antibiotic/antimycotic and either 1.5 × 10^4^ cells/mL of HEK293T or 2.5 × 10^4^ cells/mL of SH-SY5Y cells. After the 48 h transfection, the luciferase activity was quantified in the Synergy Mx Microplate Reader (Agilent, Santa Clara, CA, USA) by the Dual-Luciferase Reporter Assay System’s protocol (Promega, Fitchburg, WI, USA).

### RNA isolation and real-time RT-PCR

Total RNA was extracted from patient salivary gland tissues using the phenol-chloroform method with TRIzol reagent (Invitrogen, 15596026). The RNA was subsequently purified and isolated with the PureLink™ RNA Micro Kit (Invitrogen, #12183016), according to the manufacturer’s instructions. An average of 1500 ng of total RNA was used to synthesize first-strand cDNA (NZY First-Strand cDNA Synthesis Kit, MB125). SYBR-green quantitative PCR (CFX384 Touch™ Real-Time PCR Detection System, Bio-Rad) was performed using specific primers. Human *RACGAP1*, sense primer: CGAAGTGCTCTGGATGTTA, antisense primer: TTGCTCCTCGCTTAGTTG; Human β-actin, sense primer: ACAGAGCCTCGCCTTTGCCG, antisense primer: CACCATCACGCCCTGGTGC. The fold change in gene expression was calculated using the ΔΔCt relative expression method (Livak method) and, primers for β-actin were used as the endogenous control and calculated separately for each sample and respective condition.

### Statistical analysis

All measurements were performed with the researcher blinded to the experimental condition. Data are shown as meanLJ±LJSD. Unpaired t-tests were used for comparing differences between two groups, while one-way ANOVA followed by Sisak’s multiple comparisons was applied to identify significant differences among multiple groups. For the in vivo experiments the sample size was chosen based on previous research[13, 18]. For the *in vitro* analysis all the experiments were performed at least three times. Statistical significance was determined using the GraphPad Prism Software version 8 being significance determined by *pLJ<LJ0.05, **pLJ<LJ0.01, ***pLJ<LJ0.001 and ****pLJ<LJ0.0001. Statistical tests and sample sizes are indicated in each figure legend.

Regarding the genetic studies, since we included in the analysis several members of the same family, each patient was “nested” in his/her family. For this purpose, analyses were conducted taking into account the non-independency of AO, using generalized estimating equations (GEEs). In this model, we associate the different variants with AO (which is the dependent variable), using the most common genotype as the reference and adjusting for gender. The unstandardized coefficient (B) corresponds to the mean AO variation observed in the individuals carrying a specific genotype when compared with the reference category. To correct for multiple testing as we studied 9 genes, we applied a Bonferroni correction (α was set at 0.006 in the GEE analysis). Statistical analyses were performed using IBM SPSS Statistics software (v.29) (IBM, Armonk, NY, USA).

## Acknowledgements

Thy1-EB3–GFP and Thy1-Mito-RFP mice were provided by Dr Thomas Misgeld (Technical University of Munich, Germany). We thank i3S Scientific Platforms (Animal Facility, Cell Culture and Genotyping, Histology and Electron Microscopy, Proteomics, Advanced Light Microscopy (PPBIPOCI-01-0145-FEDER- 022122).

This work was supported by: 2021 RD G- Transthyretin Amyloid Polyneuropathy (ATTRV-PN) research competitive grant program from PFIZER to MAL; FEDER - Fundo Europeu de Desenvolvimento Regional funds through the COMPETE 2020 – Operacional Programme for Competitiveness and Internationalisation (POCI), Portugal 2020, and by Portuguese funds through FCT - Fundação para a Ciência e a Tecnologia/Ministério da Ciência, Tecnologia e Ensino Superior in the framework of the project POCI-01- 0145-FEDER-028336 (PTDC/MED-NEU/28336/2017) to MAL; Fundação para a Ciência e a Tecnologia, 2022.01656.PTDC to CL; National Funds through FCT – Fundação para a Ciência e a Tecnologia under the project IF/00902/2015 to MAL; Ministry of Science and Technology, Taiwan Government (111-2320-B-002- 079) to S-TH; and National Taiwan University Hospital, Taipei, Taiwan (UN110-014) to S-TH. MAL is supported by CEECINST/00091/2018; MIOS by 2023.08100.CEECIND; EC by a fellowship grant UI/ BD/154392/2023; and MS by the program DL 57/2016 – Norma Transitória.

