## Supplementary Data for "Rac1 inhibition prevents axonal cytoskeleton dysfunction in Transthyretin Amyloid Polyneuropathy"

**Supplementary Figures**

**Supplementary Fig. 1**

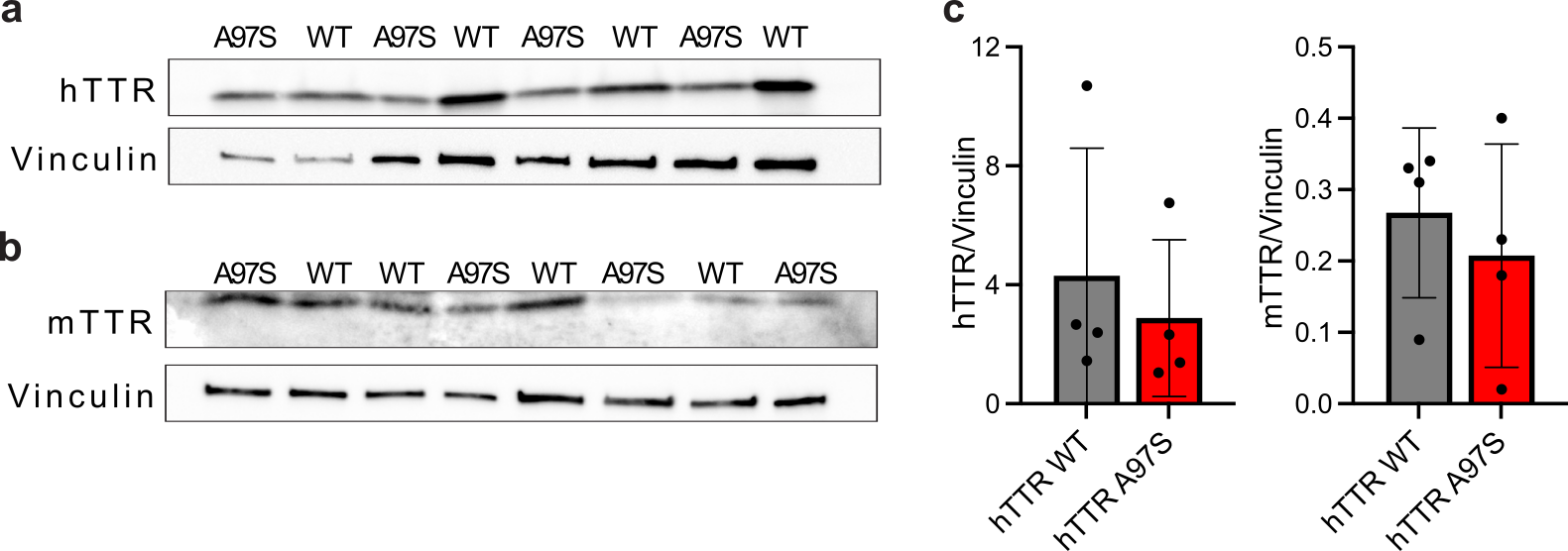

**Supplementary Fig. 1** TTR levels are similar in hTTRWT and hTTRA97S sural nerves. **a-c** Representative western blot analysis **(a, b)** and quantification **(c)** of the levels of human TTR (hTTR) **(a)** and mouse TTR (mTTR) **(b)** in 9-month-old hTTRWT and hTTRA97S sural nerves. TTR levels are relative to Vinculin. Data represent mean ± SD (n=4 animal per genotype).

**Supplementary Fig. 2**

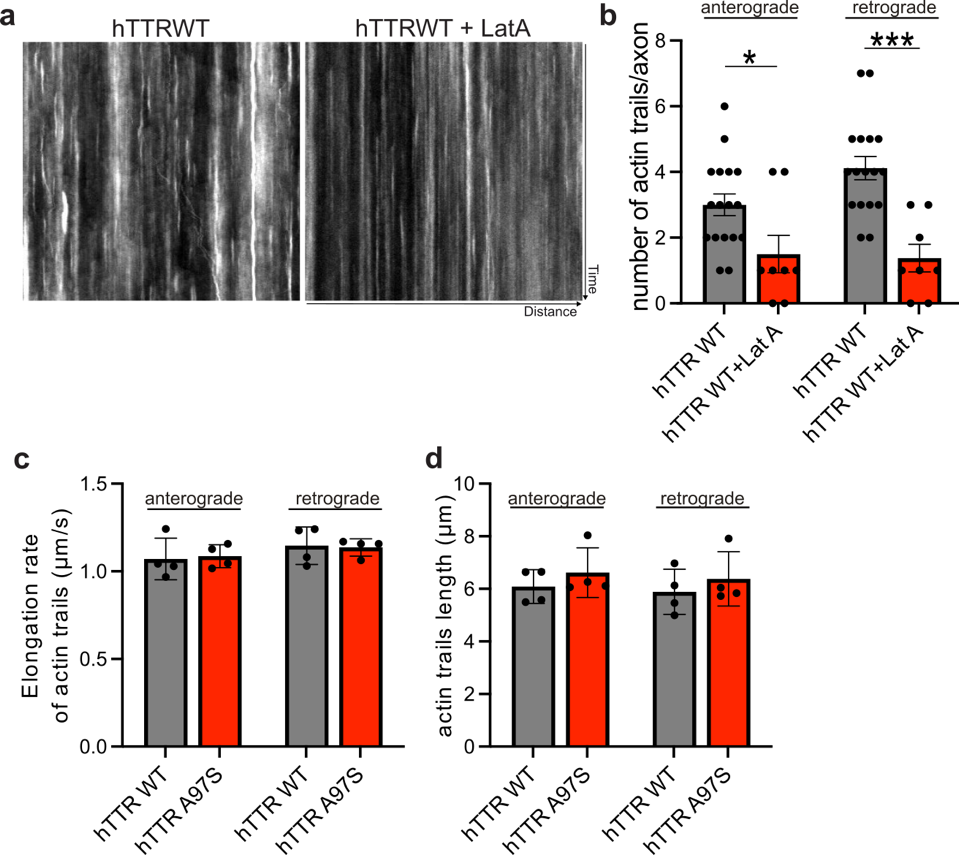

**Supplementary Fig. 2** Actin trail dynamics in DRG neurons*.* **a-b** Representative kymographs of imaged axons transfected with GFP:UTR-CH (to label F-actin) **(a)** and respective quantification of the number of retrograde and anterograde actin trails **(b)** in DIV2 hTTRWT untreated (hTTR WT) or treated with latrunculin A (hTTR WT+LatA). Data represent mean ± SD (n= 8-17 axons/conditions. Representative experiment). *p<0.05, **p<0.01 by Student’s t test. **c-d** Quantification of the rate **(c)** and length **(d)** of elongation of actin trails in DIV2 hTTRWT and hTTRA97S neurons. Results presented as mean ± SD (n= 4 independent experiments).

**Supplementary Fig. 3**

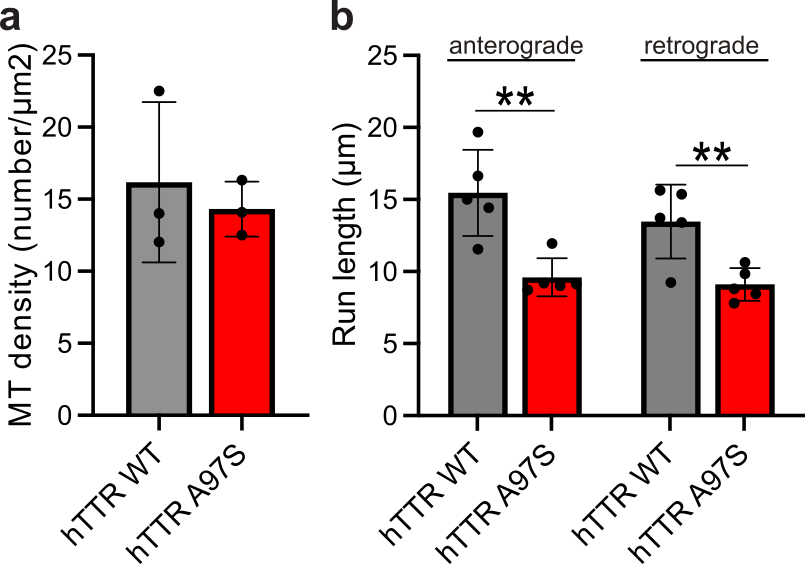

**Supplementary Fig. 3** Axonal microtubule density and mitochondrial run length in 9-month-old hTTRA97S sural nerves*.* **a** Quantification of axonal microtubule (MT) density from hTTRWT and hTTRA97S sural nerves. Results are plotted as mean ± SD (*n* = 3 animals/genotype). **b** Mitochondrial run length in hTTRWT-Thy1-MitoRFP and hTTRA97S-Thy1-MitoRFP sural nerve axons. Data represent mean ± SD (n=5 animals/genotype; 5 axons/animal).

**Supplementary Fig. 4**

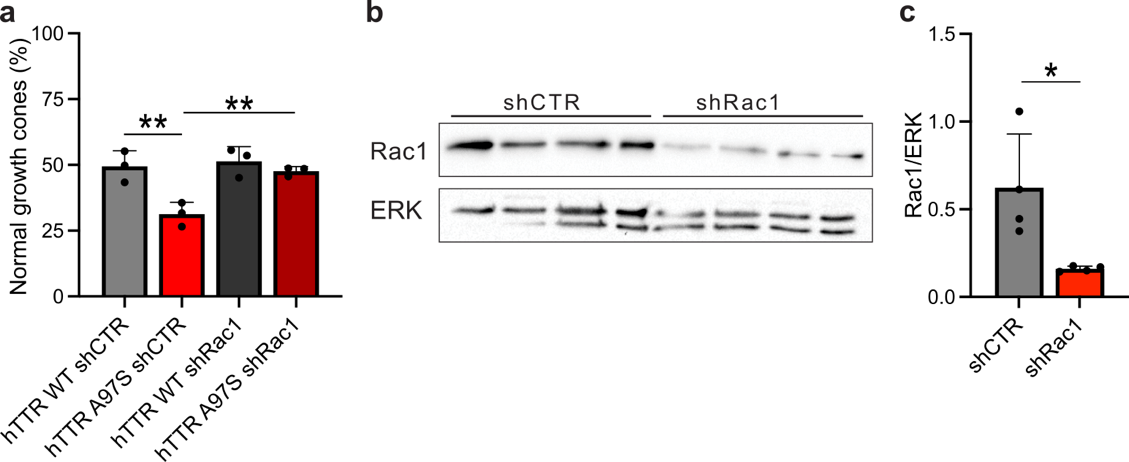

**Supplementary Fig. 4** Rac1 mediates cytoskeleton defects in hTTRA97S neurons. **a** Quantification of the percentage of normal growth cones ones in hTTRWT and hTTRA97S DRG neurons transfected with shRNA scramble (shCTR) or shRNA Rac1 (shRac1). Data represent mean ± SD (n=3 independent experiments). **p<0.01 by One-way ANOVA with Sidak’s multiple comparisons test. **b-c** Representative western blot analysis **(b)** and quantification **(c)** of the levels of Rac1 in N1E-115 cells transfected with shRNA scramble (shCTR) or shRNA Rac1 (shRac1). Rac1 levels are relative to ERK. Data represent mean ± SD (n=4 independent samples/conditions). *p<0.05 by Student’s t test.

**Supplementary Tables**

**Supplementary Table 1. List of RAC1-specific guanine nucleotide exchange factors (GEFs) and GTPase-activating proteins (GAPs).**

|  | Gene | References |
| --- | --- | --- |
| RAC1-specific GEFs | *SWAP70* | [1] |
|  | *DOCK1* | [2, 3] |
|  | *DOCK4* | [4, 5] |
|  | *DOCK5* | [6] |
|  | *DOCK6* | [7] |
| RAC1-specific GAPs | *ARAP2* | [8] |
|  | *RACGAP1* | [9, 10] |
|  | *ARHGAP31* | [11] |

**Supplementary Table 2. Quantification of F-actin dynamics in DRG neurons.**

|  | DRG neurons | | Hippocampal neurons | |
| --- | --- | --- | --- | --- |
| Direction | **Retrograde** | **Anterograde** | **Retrograde** | **Anterograde** |
| Mean length/µm | 5.889±0.4283 | 6.088±0.3203 | 8.87 ± 0.22 | 8.85 ± 0.18 |
| Rate of polymerization/$\boldsymbol{\mu}$m.s^-1^ | 1.146±0.05337 | 1.070±0.05937 | 0.99 ± 0.01 | 0.99 ± 0.01 |

Values are presented as mean ± SEM. Data from hippocampal neurons derived from Ganguly A, Tang Y, Wang L, et al. A dynamic formin-dependent deep F-actin network in axons. Journal of Cell Biology. 2015;210(3):401-417.
